# SCNIC: Sparse Correlation Network Investigation for Compositional Data

**DOI:** 10.1101/2020.11.13.380733

**Authors:** Michael Shaffer, Kumar Thurimella, John D. Sterrett, Catherine A. Lozupone

**Author notes:** Current: Department of Soil and Crop Sciences, Colorado State University, Fort Collins, CO 80523, USA. Current: Department of Chemical Engineering and Biotechnology, University of Cambridge, Cambridge, UK. These authors contributed equally to this work (Michael Shaffer, Kumar Thurimella).

## Abstract

**Background:** Microbiome studies are often limited by a lack of statistical power due to small sample sizes and a large number of features. This problem is exacerbated in correlative studies of multi-omic datasets. Statistical power can be increased by finding and summarizing modules of correlated observations, which is one dimensionality reduction method. Additionally, modules provide biological insight as correlated groups of microbes can have relationships among themselves.

**Results:** To address these challenges, we developed SCNIC: Sparse Cooccurrence Network Investigation for compositional data. SCNIC is open-source software that can generate correlation networks and detect and summarize modules of highly correlated features. Modules can be formed using either the Louvain Modularity Maximization (LMM) algorithm or a Shared Minimum Distance algorithm (SMD) that we newly describe here and relate to LMM using simulated data. We applied SCNIC to two published datasets and we achieved increased statistical power and identified microbes that not only differed across groups, but also correlated strongly with each other, suggesting shared environmental drivers or cooperative relationships among them.

**Conclusions:** SCNIC provides an easy way to generate correlation networks, identify modules of correlated features and summarize them for downstream statistical analysis. Although SCNIC was designed considering properties of microbiome data, such as compositionality and sparsity, it can be applied to a variety of data types including metabolomics data and used to integrate multiple data types. SCNIC allows for the identification of functional microbial relationships at scale while increasing statistical power through feature reduction.

## BACKGROUND

Microbial communities play important roles in environmental and human health systems and can often reach great complexity. In these rich ecosystems, microbes interact with each other, forming relationships based on predator-prey dynamics [1], competition for resources [2], cross-feeding of small compounds, [3] and other factors. Identifying correlated pairs of microbes can suggest potential interactions or shared environmental preferences. Accordingly, studies have identified complex networks of co-occurring microbes in a variety of different environments ranging from the human mouth and gut [4] to soil [5] and stream ecosystems [6].

To detect correlations between microbes, a variety of methods have been developed. While traditional correlation metrics are used by some [7–9], newer methods have been developed that take into account the properties of 16S rRNA sequencing data [10–12]. A recent review tested these methods on a variety of models and identified some methods that performed better than others in ways that can depend on underlying data characteristics [13]. Although these tools are useful for finding pairwise relationships between organisms, less attention has been given toward developing methods for finding correlations among groups of microbes.

One way to explore complex interactions is to form networks in which correlated organisms are joined with an edge, and highly correlated sets of microbes are defined. Here, we refer to these sets as modules, which are synonymous to clusters or groups. There are two primary benefits of finding modules of correlated microbes. First, the combination of microbes in a module could be further explored to understand microbial interactions, such as cross-feeding relationships, or shared environmental niches [5, 14–16]. Second, considering correlation structure among microbes can aid in statistical analysis aimed at uncovering relationships between microbes and other environmental factors. Specifically, by eliminating or summarizing highly correlated features, dependence between features is decreased. Feature reduction will increase accuracy of methods that assume the independence of features such as false discovery rate technique (FDR) measurements like the Benjamini-Hochberg Correction [17], and statistical power is increased by reducing the number of feature comparisons.

One workflow for considering groups of correlated microbes in downstream statistical analyses requires three steps: first, correlations between microbes must be measured and used to form a network; second, modules must be identified; and third, abundance of the microbes in modules must be summarized for use in subsequent statistical analyses. One software tool that has implemented this workflow, developed for application to gene expression data, is weighted gene correlation network analysis (WGCNA)[18]. WGCNA builds correlation networks based on a correlation coefficient (such as Pearson, Spearman, or biweight midcorrelation [19]), and detects modules as subtrees in a hierarchical cluster of features [20]. Modules are summarized by setting module abundance to that of network hubs or an eigenvector of the abundance of all module members [18].

Several groups have used WGCNA to find correlations within 16S rRNA sequencing data [21–24], but this approach may not be appropriate for several reasons [25]. First, the correlation metrics implemented in WGCNA do not account for sparsity and compositionality. Most sequencing-based microbiome datasets are sparse (i.e. there are many zeros) and compositional, meaning they only carry information on relative abundances of taxa instead of absolute abundances, which can lead to the detection of spurious correlations if proper statistical methods are not used [26]. Thus, the use of WGCNA for compositional data may be leading to the detection of spurious edges in microbiome networks. Second, the primary method WGCNA uses to pick modules assumes the correlation network will have a scale-free topology that may not be relevant to microbiome data [27]. Third, summarizing modules through identifying hub taxa works well in gene expression where a single transcription factor can control the expression of many genes, but may not be appropriate in microbial communities. Both the hub and eigenvector approaches to module summarization do not allow for output tables that maintain the total counts of microbial abundance per sample. Therefore, the hub and eigenvector approaches cannot be used with tools developed for microbiome data analysis that make assumptions based on total sample counts, such as ANCOM [28] or metagenomeSeq [29].

Optimal methods for identifying and summarizing modules of correlated features in 16S rRNA sequencing data have not been deeply explored. One study [25] recommended an ensemble approach for correlation detection, and the Louvain modularity maximization (LMM) method [30] to identify modules. LULU is a tool that follows a binning approach towards OTUs that co-occur, but only does so if they are highly phylogenetically related [31]. Another tool, CoNet, uses an ensemble approach to build and visualize networks [32]. However, no implementation of module summarization was made available for downstream statistical analysis.

To address these gaps, we have developed a tool for sparse, compositional correlation network investigation for compositional data (SCNIC), which uses methods optimized for microbiome data analysis. SCNIC is available as standalone Python software, via Bioconda [33] and the package installer for Python (pip), and as a QIIME 2 plugin [34]. The source code for SCNIC and the QIIME 2 plugin is freely available on GitHub (https://github.com/lozuponelab/SCNIC, https://github.com/lozuponelab/q2-SCNIC) under the BSD-3-Clause License.

## MATERIALS AND METHODS

### The SCNIC method

SCNIC takes a feature table containing counts of each feature in all samples as input and performs three steps: 1) a correlation network is built, 2) modules are detected in the network and 3) feature counts within a module are summed into a new single feature (identified as “module-*x*” where x is whole numbered consecutively starting at zero)(Figure 1). The modules are ordered based on size, where the lower numbered modules have a larger number of members compared to higher numbered modules. To summarize modules, SCNIC uses a sum of count data from all features in a module. There is no maximum or minimum size constraint on module size when modules are created. The newly generated modules are included in a new feature table alongside all features not grouped into a module. This maintains the total counts per sample, allowing for downstream analyses with tools that have assumptions related to total sample counts. SCNIC produces a graph modeling language (GML) format [35] file compatible with Cytoscape [36] for network visualization in which the edges in the correlation network represent the positive correlations which are stronger than a user specified R-value cutoff (between 0 and 1), a file describing which features compose each defined module, and a feature table in the Biological Observation Matrix (BIOM) [37] (Figure 1).

**Figure 1:**
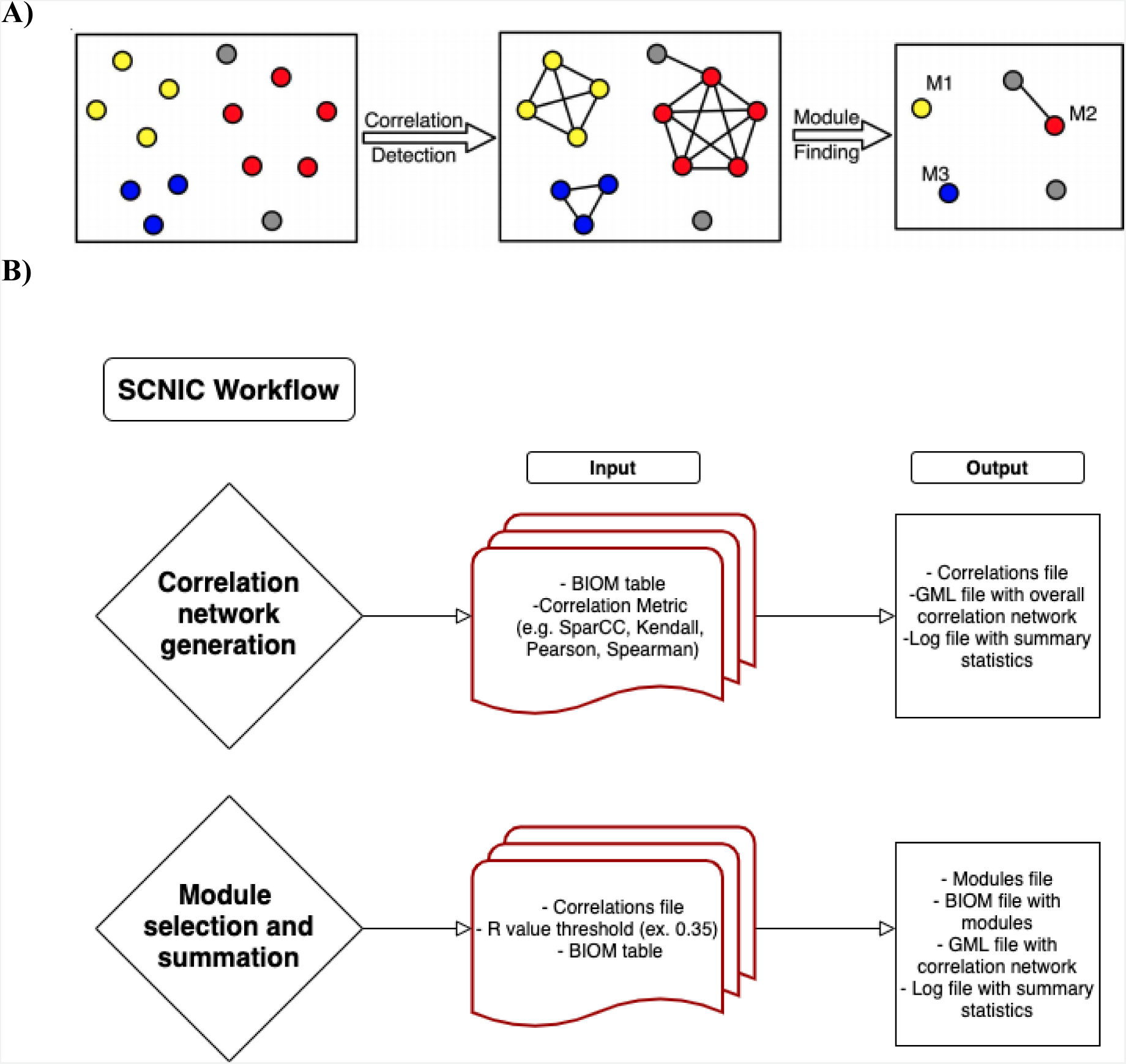
SCNIC Schematic and Data Flow. A. The basic process of SCNIC involves first identifying pairwise correlations between species and using them to build a correlation network. Modules of correlated features are identified and then summarized for downstream statistical analysis, or multi-omic analysis between modules of microbes and other feature types. B. The input to SCNIC comes in the form of a count table in BIOM format. The first step takes the table and generates a correlation table and network. The table is in a tab delimited format and the network is in GML format and can be used to visualize the network in Cytoscape. Modules are detected and summarized in the final step which generates a module membership file indicating which features are in each module. The collapsed BIOM table contains the same total counts per sample as the original table, but with less features. All features not included in modules are retained with their original counts and all modules have a total count per sample of the sum of the counts of all features in that module.

SCNIC allows users to choose between multiple methods for detecting correlations and of defining modules of co-occurring microbes. For correlations, SCNIC can implement traditional correlation metrics (including Pearson’s *r*, Spearman’s *⍴* and Kendall’s τ) or the compositionality- and sparsity-aware correlation metric from SparCC [38, 39] to correct for aspects of microbiome data. SparCC has been shown to perform well in detecting correlations compared to other correlation measures [13]. Specifically, SparCC performs well in communities with an inverse Simpson index above 13 (which would be indicative of a high number of successful species, a complex food web, and many ecological niches, as would be seen in many high biomass microbial communities such as gut or soil microbiomes) [39, 40], and it thus was chosen as the default metric.

To define modules of co-correlated features, we implement two methods: 1) Louvain modularity maximization (LMM) and 2) a novel shared minimum distance (SMD) module detection algorithm; unlike WGCNA, neither of these algorithms make assumptions about network topology. LMM was previously proposed as a method for clustering correlation networks of microbes into modules [30]. LMM works by first assigning one module per feature. Each pair of adjacent modules are joined and the change in modularity (defined by the number of edges within the module compared to outside) is calculated for each module. The pair which increases the mean modularity of the network the most is then joined. This process is repeated until the modularity of the network is not increased. LMM uses two parameters provided by the user: The first parameter, R-value, defines the minimum correlation coefficient for defining an edge between features. The second parameter, gamma (also referred to as resolution), controls the size of modules detected, with large gamma values yielding larger modules.

WGCNA and LMM have a potential weakness in that modules can contain pairs of taxa that are not strongly correlated (e.g. if they are several steps away from each other in the network). To address this weakness, we also implement the SMD method to ensure that correlations between all pairs of features in the module have an R-value greater than the user provided minimum (Figure 2). Specifically, the SMD method defines modules by first applying complete linkage hierarchical clustering to correlation coefficients to make a tree of features. Next, SMD defines modules as subtrees where correlations between all pairs of tips have an R-value above the specified value. SMD has been set as the default method in SCNIC because of the desirable property of only producing modules where all features are correlated over a user-specified threshold.

**Figure 2:**
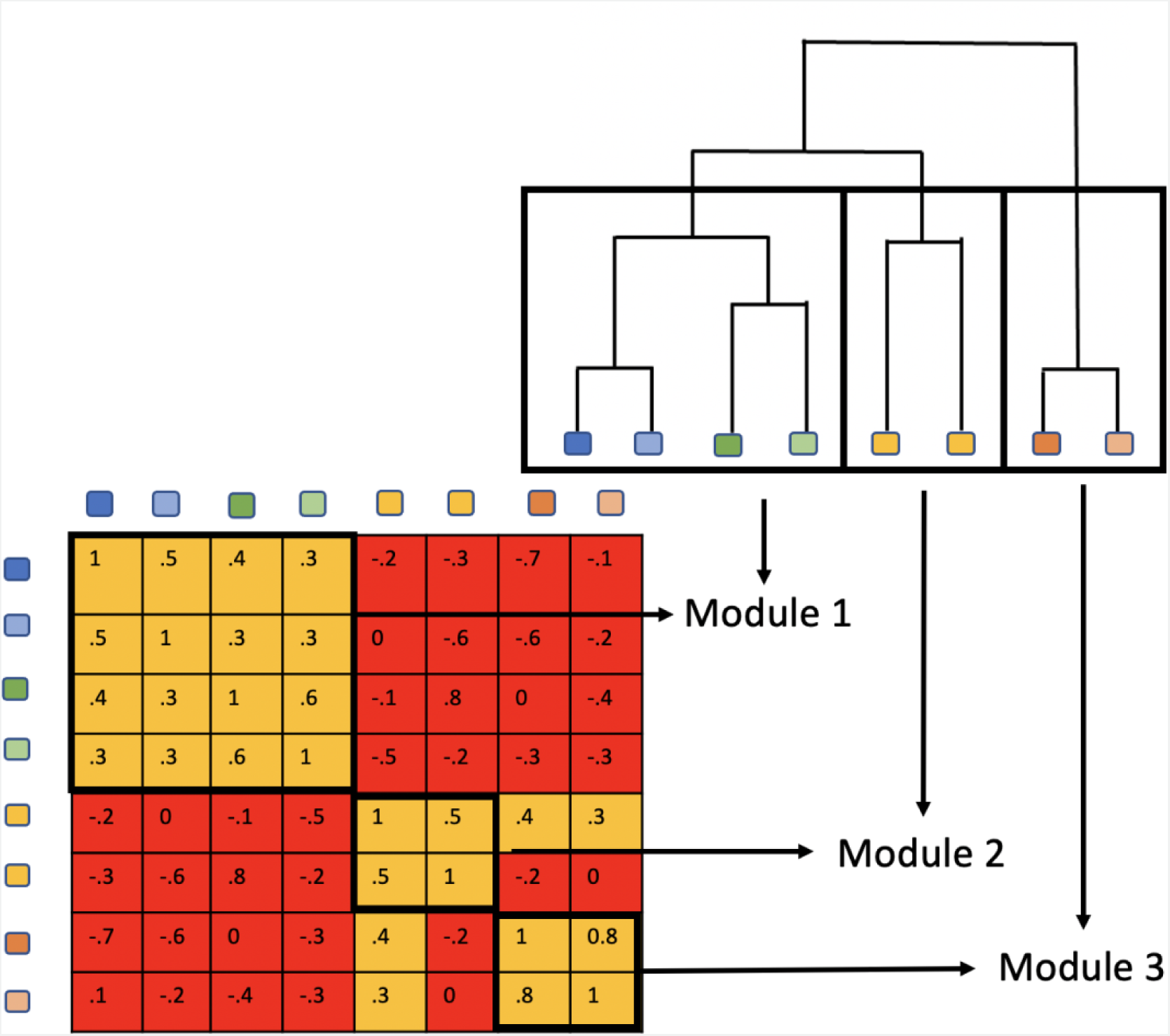
SMD Algorithm for Defining Modules. SMD defines a module as a group where all features have a correlation above a given threshold. To do this SMD first uses complete linkage hierarchical clustering on correlation coefficients to create a tree. Each module is defined as a subtree where the correlation coefficients between all tips are greater than the threshold.

A large proportion of microbiome studies sample highly uneven communities which leads to strong compositionality-driven artifacts [26, 40, 41]. Because of this, we use SparCC, specifically the implementation of FastSpar [39], as the default correlation measure. SparCC was used as the correlation metric based on analysis that suggested a high precision in the number of correct edges recovered when correlations were calculated in synthetic data [13]. SCNIC additionally includes the option of using Pearson’s *r*, Spearman’s *⍴* and Kendall’s τ to evaluate non-compositional or dense data types.

### Evaluating the SMD algorithm using simulated data

Since SMD has not been applied to microbiome module detection before, we compared SMD to LMM using simulated data. In order to evaluate the performance of SMD for module detection under different parameter settings and compare it to LMM, we simulated a wide range of networks. The simulations had networks with similar characteristics to those seen in networks generated from microbiome datasets. These included networks with power law degree distributions (N=175) with values of ɑ, the exponent term of the power law formula (*y* = *kX^-ɑ^*), varying between 1.8 and 2.6, as well as networks with regular degree distributions (N=200) with *p*, the probability of one node being connected to another, varying from 0.001 to 0.2. The power law and regular degree distribution networks were created using the NetworkX v2.6.3 implementations of configuration_model and erdos_renyi_graph, respectively, and all had a size of 500. The networks with power law degree distributions had modularity values between 0.2 and 0.9, with higher ɑ corresponding to higher modularity, and the networks with regular degree distributions had modularity values between 0.07 and 0.98, with lower *p* corresponding to higher modularity. Higher modularity scores indicate many connections within modules and fewer connections between modules. We then calculated SMD and LMM partitions (with LMM gamma = 1) of each network and compared the homogeneity between the two partitions. Because SMD modules are smaller than LMM modules, we used the homogeneity metric described by Rosenberg and Hirschberg [42] (implemented via Scikit-learn v0.24.2) to assess whether nodes partitioned together by SMD are a subset of the module partitioned by LMM. A score of 1 represents that all nodes in SMD modules represent sub-modules of LMM-partitioned modules, whereas a score of 0 represents that no two nodes that were classified by SMD into the same module were partitioned into a module together by the LMM method.

### Demonstrating the use of SCNIC

We demonstrate the use of SCNIC with two example datasets. These are 1) a study that used 16S rRNA sequencing of fecal material to compare microbiome composition in individuals with and without HIV and in men who have sex with men (MSM) who were at a high risk of contracting HIV [43], and 2) a dataset analyzing the microbiome of water samples at various depths in two of the Great Lakes. We chose these two datasets so that we could evaluate performance using datasets from both host-associated and free-living microbiomes. We also used the Great Lakes dataset to compare module size and modularity between SMD and LMM selected modules.

#### HIV dataset

The HIV data set was retrieved from NCBI SRA accession number SRP068240, and samples from the BCN0 cohort were used for these analyses. Reads were error corrected, quality trimmed, and primers were removed using default parameters in BBTools [44]. DADA2 [45] was used to define amplicon sequence variants (ASVs) with reads trimmed from the left by 30 base pairs and truncated at 269. ASVs were binned into operational taxonomic units (OTUs) using USEARCH [46] at 99% identity using QIIME 1 [47]. A phylogenetic tree was made using a single representative sequence from each OTU and the SEPP protocol [48, 49] using QIIME 2 [34]. We evaluated the average phylogenetic distance between OTUs in the same module using the *distance* method of Biopython [50, 51]. Taxonomy was assigned using the Naive Bayes QIIME 2 feature classifier, version gg-13-8-99-515-806-nb-classifier.qza.

The original study describing these data showed a strong divergence in gut microbiome composition in MSM compared to non-MSM independent of HIV infection status and more subtle differences associated with HIV infection when controlling for MSM behavior. The goal of our analysis was to evaluate whether comparing gut microbiome composition between HIV negative MSM and non-MSM with SCNIC modules provide additional significant taxa compared to without, and additional insights as to which taxa that differ with MSM also are in turn demonstrating co-correlated structure with each other. Co-correlation of microbes may indicate that they are a part of a broader community type, interact with each other, or have shared environmental drivers of their prevalence. A further goal of this analysis is to examine the effects of using different R-value thresholds on the results. The SMD method was specifically used with SparCC R-value thresholds between 0.20 and 1.0, with 0.05 increments.

#### Great Lakes dataset

The Great Lakes dataset was previously published as part of the Earth Microbiome Project [52]. This study evaluated patterns of microbial relative abundance across depths in Lake Michigan (N=16) and Lake Superior (N=33), with depth of samples collected ranging from 5 to 3654 meters. The study additionally recorded data on pH and salinity. The Great Lakes data set was retrieved from QIITA accession number 1041 [53]. ASVs were found using DADA2 with a left trim of 30 and a truncation length of 135. OTUs were subsequently picked on the ASVs using VSEARCH [54] with a 99% identity threshold, resulting in 3,871 OTUs. These steps were done with QIIME 2 [34]. SCNIC was applied with the SMD method and .2, .4 and .65 R-value thresholds.

#### Comparison of SMD to LMM using the Great Lakes dataset

To identify differences in module structure from SMD versus LMM partitions, we assessed the module size and modularity of 221 separately partitioned networks from the Great Lakes dataset using varying parameters for SCNIC. The parameters included SCNIC R thresholds ranging from 0.1 to 0.7 and gamma ranging from 0.15 to 0.9 for LMM.

### Evaluating effects of applying SCNIC to discern microbes that differ between groups in the HIV and Great Lakes datasets

OTUs/modules that differed with MSM status (HIV study) and between lakes (Great Lakes study) were identified using ANCOM [28] for each feature. For the first study, we focused on evaluating differences in the microbiome between MSM and non-MSM without confounding by HIV infection status, by only using samples from HIV negative individuals. We chose ANCOM because it is also a tool designed specifically for working with compositional microbiome data. ANCOM was applied to the original feature table where SCNIC was not applied, as well as to feature tables output from SCNIC using SparCC at different R-value thresholds with the SMD algorithm.

While applying SparCC, SCNIC uses the recommended practice of the SparCC manuscript of filtering based on average relative abundance across samples [38]. The SparCC manuscript suggests this filter because removing features with high abundances, even in a few samples, will upset the ability of the method to control for the number of reads per sample in its compositionality adjustment. Because this method can retain OTUs that are highly abundant in only a single sample, we removed features that had 0 values in more than 5% (∼ 29/146) of samples before applying ANCOM but after applying SparCC. Significant differences between groups were determined as those above the W-value threshold determined by ANCOM.

### Time and memory usage by SCNIC

To evaluate the memory resources needed by SCNIC, we ran the SCNIC modules step locally on a 2015 MacBook Pro with 16 GB RAM with a 2.5 GHz Quad-Core Intel Core i7 processor for both the Great Lakes dataset and an integrated microbiome-metabolome dataset with 1,301 features, which will be referred to as the GT dataset. The runtime was recorded across 3 runs per method (SMD vs LMM) for each dataset using GNU Time, and memory was profiled using memory-profiler 0.60.0. The “within” step, which calculates correlations between features and creates the correlation network was not tested because it depends greatly on the correlation metric used, and the runtime and memory usage of FastSpar (likely the most computationally intensive correlation metric to be used in this step) have already been profiled[39]. The modules step only utilizes a correlation matrix and as such does not scale with the number of samples, only the number of features, except when the values of the count table are being summed, which is a generally trivial calculation compared to the module generation step.

## RESULTS

### Comparison of LMM to SMD on real and simulated data

In order to evaluate the relationship between modules detected with SMD versus LMM, we chose modules on the Great Lakes dataset using SMD at R-value thresholds ranging from 0.05 to 0.7 and with LMM at the same R-value thresholds and gamma values ranging from 0.15 to 0.9. We found that with both LMM and SMD, modularity increased with increasing R-value thresholds. However, SMD produced less modular partitions and smaller modules than LMM, even when LMM was applied with very low values for the gamma parameter that controls module size (Figure 3 A,B).

**Figure 3:**
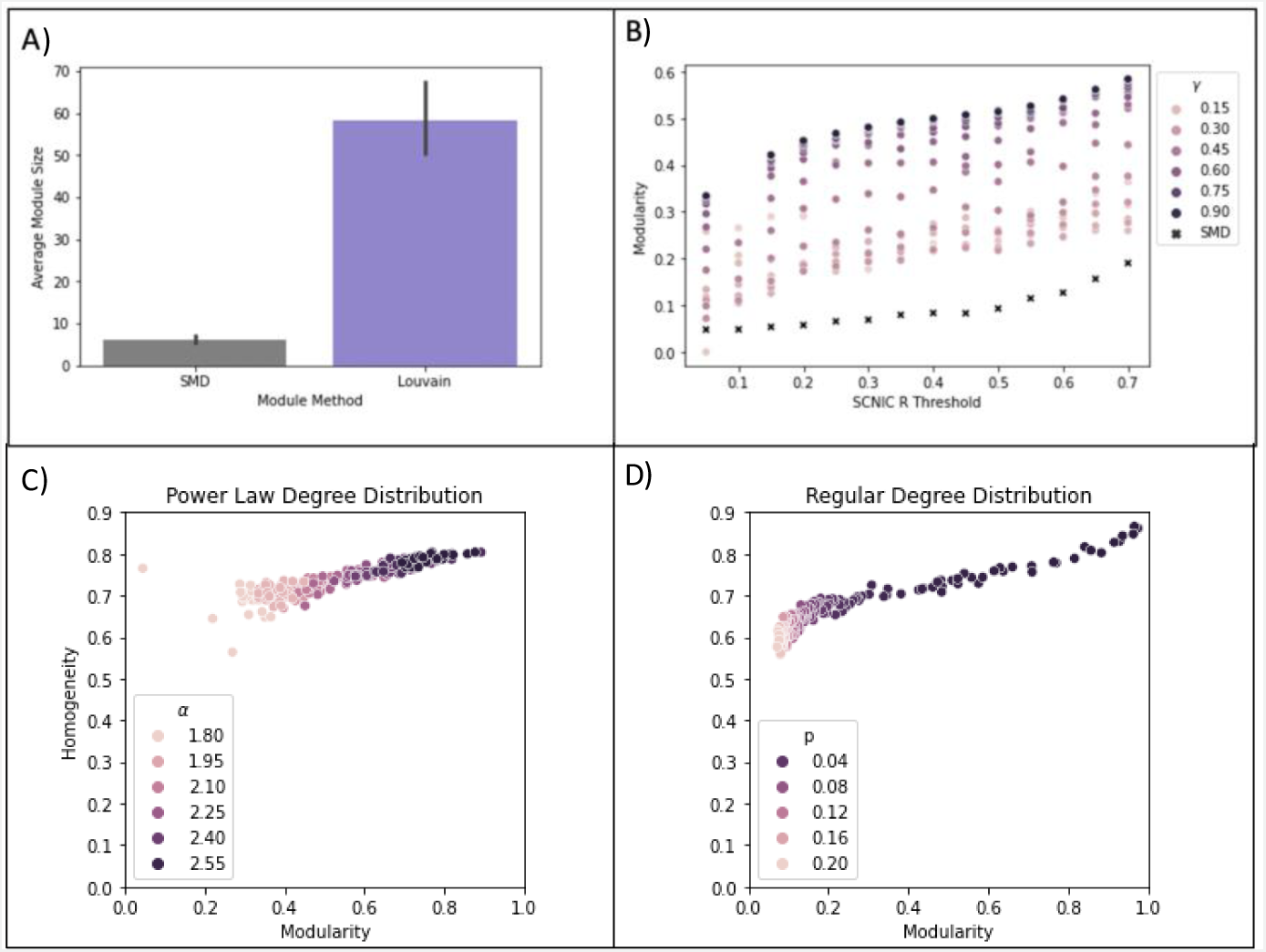
Comparison of the SMD and LMM Algorithms for Module Selection. Panels A and B show a comparison of module size and modularity between SMD and LMM selected modules on the Great Lakes data at R thresholds ranging from 0.1 to 0.7 and gamma ranging from 0.15 and 0.9 for LMM. Panel A compares average module size and Panel B shows modularity across these parameters. In Panels C and D the homogeneity of LMM versus SMD selected module was calculated using simulated networks formed using both Power-Law (Panel C) and Regular (Panel D) degree distributions. Nodes are colored by the ɑ (exponent) parameter for power law degree distributions in Panel C and by p (probability of one node being connected to any other node) in regular degree distributions in Panel D. A homogeneity of 1 denotes that all SMD modules are sub-modules of the LMM modules, whereas a homogeneity of 0 denotes that no two nodes grouped into a SMD module were partitioned into the same module by LMM.

In order to determine whether SMD produced related modules to LMM (e.g. since SMD modules are smaller, whether they represent sub-graphs of the larger LMM modules), we calculated a homogeneity score (described in methods section) between SMD and LMM modules in simulated networks. All networks contained 500 nodes. Modularity was calculated for the LMM partitions, and the homogeneity of SMD and LMM partitions was calculated. From our simulated networks, the homogeneity between SMD and LMM module partitions was between 0.55 and 0.87 (Figure 3 C,D). Notably, we found that for networks simulated with both power law and regular node degree distributions, as modularity of LMM partitions increased, the homogeneity between SMD- and LMM-partitioned modules increased (power law network Pearson R = 0.87, p < 0.001; regular network Pearson R = 0.93, p < 0.001). Thus, when network modularity is high (i.e. there is a high number of edges within the module compared to between modules), SMD partitions tend to be sub-partitions of LMM partitions.

### R-value thresholds influence module size and phylogenetic relatedness of OTUs binned into a module

A key parameter in SCNIC is the R-value threshold used to pick modules. Use of a high R-value threshold would be expected to bin only very tightly correlated microbes with strong relationships, while less stringent thresholds may identify community-level patterns representing more loosely connected microbial pairs. To illustrate this concept, we binned OTUs into modules using the SMD method at R-value thresholds between 0.2 and 1.0 using the HIV dataset. As expected, at lower R-value thresholds, more OTUs were binned into modules and lower numbers of modules of smaller average size were formed as the threshold increased (Figure 4A). To illustrate the effects of R-values thresholds on the nature of the identified modules, we compare SCNIC outputs using R-value thresholds of 0.2, 0.4, and 0.65. As shown in Figure 4, which visualizes modules in Cytoscape using SCNIC output files, the R-value threshold influences the size and connectivity of the network. We also illustrate the effect of using different thresholds by examining the correlations between OTUs that are included in the first module output by SCNIC, which is the largest module (module-*0*) (Figure 5). All OTUs in module-*0* are positively correlated with each other, since SCNIC only captures positive correlations.

**Figure 4:**
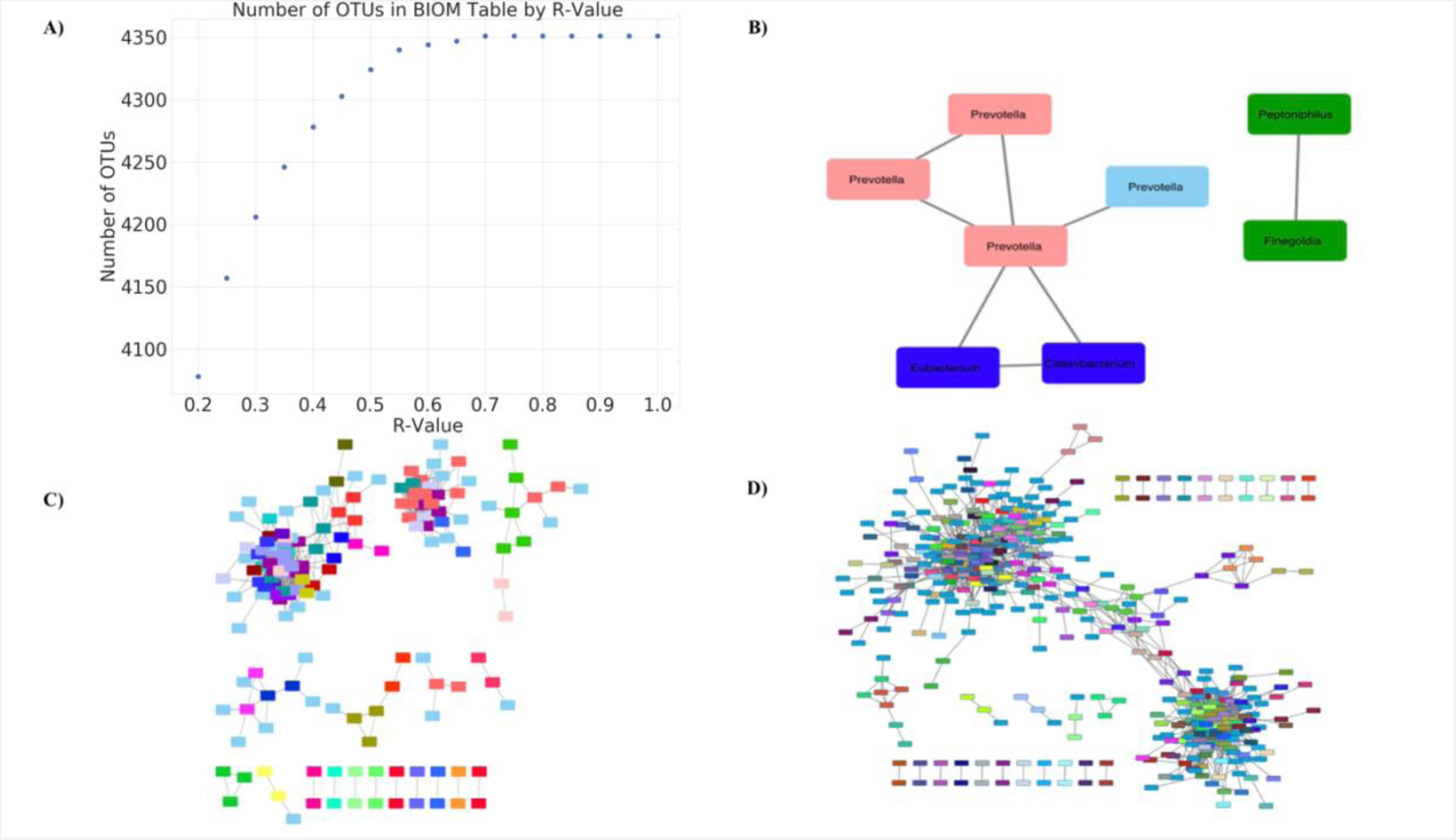
SCNIC Feature Reduction and Visualization of SCNIC Networks. A) We used SCNIC to select modules using the OTU table from the HIV dataset, the SMD module selection algorithm, and SparCC R-values ranging from 0.2 to 1.0, in increments of 0.05. The R-value is plotted against the number of features in the resulting BIOM table produced by SCNIC. As the R-value increases the number of modules decreases and the number of single features (modules + OTUs not included in modules) increases. After the R-value of 0.65, the number of features in the resulting file remained the same at 4351 features, because there were no modules that were created past a SparCC R of 0.65.The Cytoscape output allows for easy exploration and visualization of relationships between different OTUS/taxa in an interactive interface. B) R = 0.65 C) R = 0.4 D) R = 0.2. As the R-value increases, the size of the network decreases as SCNIC does not include singletons (features with no significant positive correlations) in the resulting network file. Correlation network where edges are correlations with a R-value greater than the threshold set. Nodes are OTUs and node color represents module membership (i.e. module-*0* is pink in Panel B).

**Figure 5:**
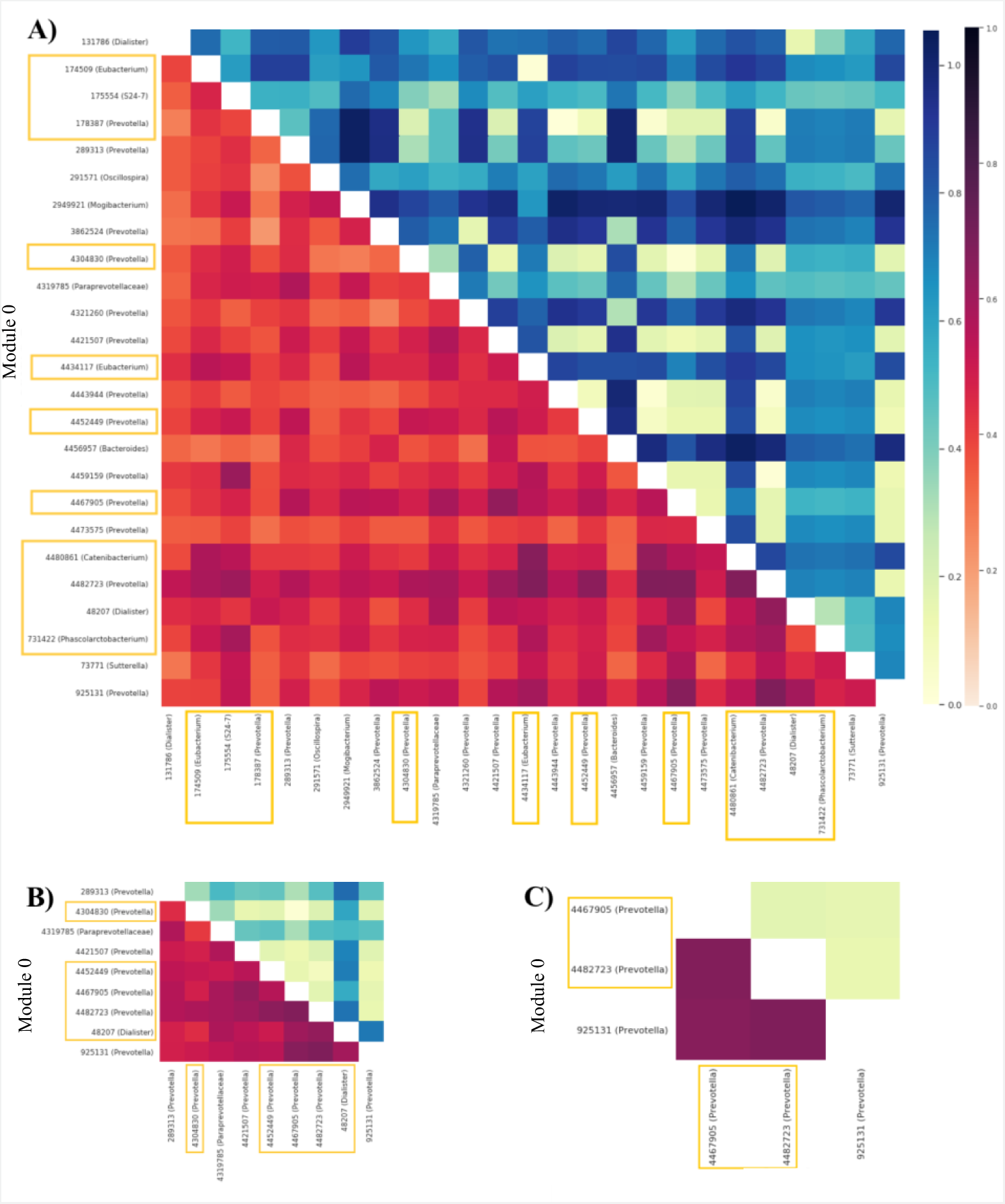
Module-*0* Across Different R-Values. Module-*0* expanded to view taxonomy and correlations amongst them at R-values of 0.2 (A), 0.4 (B), and 0.65 (C). The heatmap in the lower triangle corresponds to the correlation found by SparCC colored on a light red (low correlation) to dark red (high correlation) spectrum as defined in the color bar on the right. The heatmap in the upper triangle represents the phylogenetic distance between organism pairs colored on a yellow (small phylogenetic distance) to dark blue (high phylogenetic distance) spectrum as defined in the color bar on the right. As the R-value increases, the species in module-*0* become more phylogenetically similar. Module-*0* has 11, 5 and 2 of the significant Pre-SCNIC OTUs at R-values of 0.2, 0.4 and 0.65, and are highlighted in a yellow border.

Microbes co-occurring in the same environmental niche have previously been observed to be phylogenetically closer on average[4]. This is likely because phylogenetic relatedness has been correlated with functional relatedness, such as through having more shared genome content, leading towards success in similar environments [55]. We show that increasing the R-value threshold results in modules that contain OTUs that are more phylogenetically similar on average (Figure 5).

### Use of SCNIC influences the detection of OTUs that differ between MSM and non-MSM

We next evaluated the effects of applying SCNIC with default SparCC and SMD parameters and varying R-value thresholds on downstream statistical analysis results. To investigate differential abundance based on MSM status in the HIV dataset we used ANCOM[28]. After removing taxa that were not present in at least 5% of the samples the OTU table had 317 samples and 639 OTUs. We found that 12 OTUs were significantly different between MSM and non-MSM without using SCNIC. Using SCNIC at R-values of 0.2, 0.4, and 0.65 and running ANCOM on the filtered output feature table, we found that most of the significant features were modules at an R-value of 0.2 and 0.4 but not 0.65 (e.g. 14 of the 15 significant features were modules at R=0.2) (Table 1). This was the case even though the vast majority of OTUs were not a part of modules at the 0.4 R-value threshold (Figure 4A). The majority of 12 of the OTUs that were significant without running SCNIC, were grouped into modules with each other and with OTUs that were not individually significant without running SCNIC. These significant modules contained 74, 26, and 1 new OTU at R-values of 0.2, 0.4 and 0.65 respectively. Using SCNIC also resulted in the identification of 1, 5 and 25 (at R-values of 0.2, 0.4 and 0.65) OTUs that were individually significant that were not significant without running SCNIC, with no OTUs that were individually significant losing significance because they were binned in a module, indicating an increase in statistical power resulting from running a test like ANCOM that controls the FDR.

**Table 1:** Significant SCNIC Modules and Features Across R-Values in the HIV and Great.

| R-Value | New OTUs in Sig. Modules | New Significant OTUs | Lost Significant OTUs | # of Significant Modules | Total Significant Features |
| --- | --- | --- | --- | --- | --- |
| 0.2 | 74 | 1 | 0 | 14 | 15 |
| 0.4 | 26 | 5 | 0 | 11 | 17 |
| 0.65 | 1 | 25 | 0 | 2 | 35 |

**Table 1: Significant SCNIC Modules and Features Across R-Values in the HIV and Great** **Lakes Datasets**
| R-Value | New OTUs in Sig. Modules | New Significant OTUs | Lost Significant OTUs | # of Significant Modules | Total Significant Features |
| --- | --- | --- | --- | --- | --- |
| 0.2 | 139 | 0 | 64 | 1 | 25 |
| 0.4 | 64 | 3 | 14 | 29 | 33 |
| 0.65 | 12 | 13 | 6 | 24 | 74 |

Considering correlation structure of significant features can help in understanding the broader community context of bacteria that differ with MSM status. In module-*0* for each of the R-values, which significantly differed by MSM status in all cases, *Prevotella* was the dominant genus (Figure 5). At an R-value of 0.65, all OTUs in module-*0* were assigned to the genus *Prevotella* (Figure 5C). However, at an R-value of 0.4 module-*0* included seven *Prevotella* OTUs, one *Dialister*, and an unidentified member of the *Paraprevotellaceae* family. At the R-value of 0.2, *Prevotella* accounted for 13 of the 25 OTUs and 11 of the 12 pre-SCNIC significant OTUs were all found in this module. This suggests that individual OTUs that differ with MSM status may in some cases be a part of a consortium of diverse members that collectively display features that may contribute to differences in microbiome function.

To further explore this concept, we investigated the results generated with an R-value of 0.4, as the significant features maintain a strong level of correlation while being phylogenetically diverse. When running ANCOM on this feature table, we found that these individually significant OTUs tended to be joined into modules with other highly co-correlated microbes and that these modules significantly differed with MSM (Figure 6). Of particular note, we observe that the modules and taxa that are significantly related to MSM do not all correlate with each other. At the R-value of 0.4, module-*36* contains two taxa, *Erysipelotrichaceae* and *Clostridium* that are negatively correlated with the other significant taxa and modules (Figure 6). Module-*2* contains *Eubacterium, Catenibacterium* and *Prevotella* which are phylogenetically heterogenous but mutually co-occurring. A follow up experiment, which leverages insights that SCNIC generates, may combine different strains of microbes to assemble a community type to test for functional correlates of disease.

**Figure 6:**
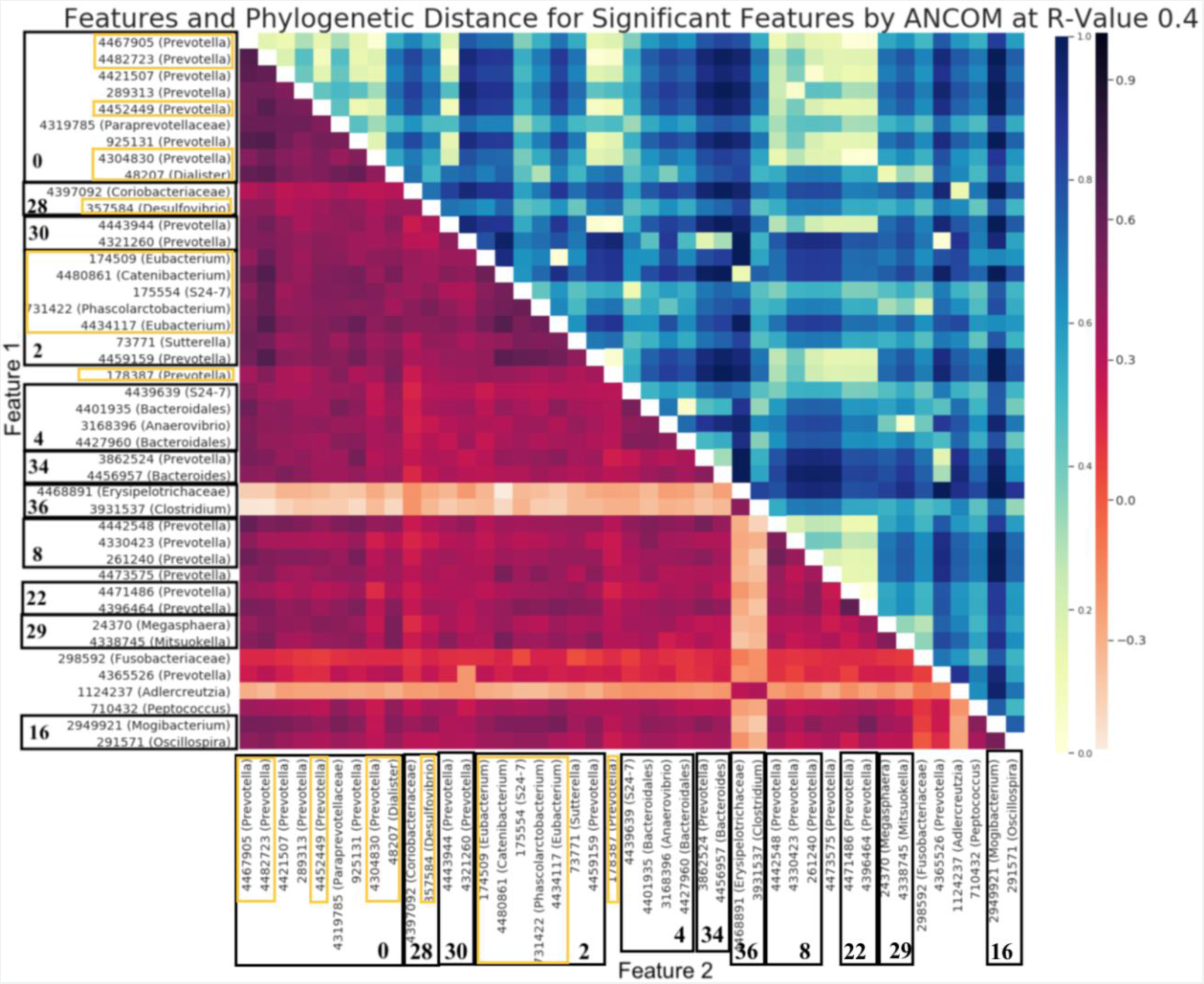
All Significant Features at R-value 0.4 Found by ANCOM. Each of the borders in the y-axis represents the different modules, with the module number bolded. The Pre-SCNIC OTUs that were significant are highlighted in a yellow border. The heatmap in the lower triangle corresponds to the correlation found by SparCC colored on a light red (low correlation) to dark red (high correlation) spectrum as defined in the color bar on the right. The heatmap in the upper triangle represents the phylogenetic distance between organism pairs colored on a yellow (small phylogenetic distance) to dark blue (high phylogenetic distance) spectrum as defined in the color bar on the right.

### Use of SCNIC results in the detection of lake associated taxa

To test consistency in patterns across different datasets, we also tested the effects of applying SCNIC with default SparCC and SMD parameters and varying R-value thresholds on results from the Great Lakes dataset. Specifically, we identified features that significantly differed between Lake Michigan (N=16) and Lake Superior (N=33) using ANCOM[28].

We began with a table of 3,871 OTUs, and 725 of these remained after removing OTUs not present in at least 5% of the samples. We found that 168 OTUs were significantly different between lakes without using SCNIC using ANCOM. Using SCNIC at R-values of 0.2, 0.4, and 0.65 and running ANCOM on the filtered output OTU table, we found that most significant features were modules at an R threshold of 0.4 but not 0.2 or 0.65 (Table 1). Use of SCNIC resulted in the detection of individual OTUs that were now significant that were not before (3 and 13 for R-value thresholds of 0.4 and 0.65 respectively). Application of SCNIC also identified many additional OTUs that become of interest because they were now part of significant modules (139, 64, and 12 OTUs at 0.2, 0.4, and 0.65 respectively; Table 1). However, unlike for the HIV dataset, several OTUs that were individually significant were no longer significant with ANCOM after applying SCNIC and this effect was the most pronounced with lower R-value thresholds (64, 14, and 6 OTUs that were significant with SCNIC were no longer significant after applying SCNIC at 0.2, 0.4, and 0.65 R-value thresholds respectively)(Table 1). This is likely because microbes that differed between lakes were binned with loosely correlated microbes that did not, leading to a loss of signal. Thus, in this case, only SCNIC with a moderate to high R-value threshold appeared to balance the benefit of the increased power and disadvantages of loss of signal from binning loosely correlated features.

### Time and memory resources used by SCNIC

SCNIC’s module generation step can be run locally on a laptop computer. For both the Great Lakes and GT datasets, which had 764 and 1,301 features respectively, SCNIC’s module generation step ran in less than 2 minutes when using the SMD method and less than 30 seconds when using the Louvain method. The Great Lakes dataset used less than 200 mebibytes (MiB) memory, and the GT dataset required a maximum of 300 MiB memory. Supplemental table 1 shows time and memory usage during SCNIC.

## DISCUSSION

SCNIC provides a method to measure correlations, find and visualize modules of correlated features, and summarize modules by summing their counts for use in downstream statistical analysis as one method for dimensionality reduction. Using SCNIC with the SMD algorithm for module detection aids in feature reduction in 16S rRNA sequencing data while ensuring a minimum strength of association within modules. As expected, our workflow identified modules in which OTUs tended to be phylogenetically related, especially at relatively high values of R. Using SCNIC, we overall achieved increased statistical power from performing less comparisons, but use of low R-value thresholds had the potential to lead to loss of significance by binning loosely correlated features. In this analysis, we used OTUs as features; however, other microbiome features can be used with SCNIC, such as ASVs, genera, or species defined with a taxonomic classifier, as well as other data types such as metabolome data. SCNIC has also been used in previously published work to perform feature reduction prior to random forest analysis with the microbiome and diverse other data types [56].

SCNIC complements existing methods because these either: 1) form correlation networks of microbes for visualization but do not have functionality for selecting and summarizing modules for downstream statistical analysis[32], 2) can select and summarize modules for downstream statistical analysis but are designed for gene expression and not microbiome data[18], only summarize features if they are phylogenetically related[31], or suggest methods for finding modules of correlated microbes but do not provide a convenient implementation[30]. SCNIC is available both as a stand-alone application and as a QIIME 2 plugin for easy integration with existing microbiome workflows.

SCNIC implements both the LMM algorithm, which had been previously recommended for selecting modules of correlated microbes [25, 57], and a novel SMD algorithm. The advantage of the SMD algorithm is that all pairs of features in the module have an R-value greater than the user-provided minimum threshold. Using real and simulated data, we showed that SMD produced smaller modules that generally represent sub-graphs of the larger LMM modules. Since the use of lower R-value thresholds similarly produced larger modules including more weakly correlated modules, we speculate that use of LMM might result in a similar trend of identifying more OTUs within significant modules, but with the disadvantage of individually significant OTUs being lost because they are combined with loosely correlated microbes that are not related to the outcome being tested.

We illustrate here that varying the R-value threshold input by the user has a great impact on the results. However, we have avoided giving specific R-value threshold recommendations here, because optimal R-values may vary across datasets and data types. Using higher R-values thresholds was more likely to identify highly phylogenetically related microbes that likely share overlapping functionality, and in principle could also identify diverse organisms with overlapping niches or highly complementary metabolic functions. Using a lower R-value threshold bins a broader community of more loosely correlated features with the risk of bringing together features which should not be grouped and loosing significance of OTUs – as was illustrated in the Great Lakes dataset analysis conducted here. By summarizing correlated features, SCNIC mitigates overcorrection in multiple test adjustments by reducing the number of taxa and false discovery rate for downstream analysis. When these organisms are grouped into a broader module that is truly independent from other modules, any penalties on two highly similar features may be avoided in statistical analysis.

The results of our HIV dataset analysis confirm original findings, as well as those of another study[58], but included many new significantly associated taxa. SCNIC also assists in interpretation of microbiome data by identifying correlations among these taxa. Our results recapitulated those of the original publication of these data and previous HIV microbiome studies that all found enrichment of *Prevotella* with MSM status [43, 58–60]. However, our analyses provide additional insight by identifying correlations between differentiating taxa. For instance, in module-*0*, which was more abundant in MSM samples, OTUs assigned taxonomically to the *Prevotella* genus are correlated with two OTUs identified as *Eubacterium biforme* (which has recently been renamed *Holdemanella biformis* [61]). *Prevotella copri* has previously been associated with increased inflammation [59] while *in vitro* stimulations of human immune cells have found that *P. copri* did not induce particularly high levels of inflammation but *E. biforme* did [60]. This strong correlation between *P. copri* and *E. biforme* in MSM could explain the increased inflammation seen in individuals with higher levels of *P. copri,* with *E. biforme* being the true driver. Indeed, MSM status has previously been associated with increased inflammation [62, 63]. With the use of SCNIC, this correlation highlighted a route of mechanistic understanding which could be functionally followed up on in further experimental studies.

SCNIC detected multiple significant modules, of which none of the OTUs within were significant when analyzed separately. Module-*20*, which was associated with MSM status, is the fourth most significant feature at R-value of 0.2, and is made up of *Acidaminococcus*, *Megasphaera*, and *Mitsuokella* species. These are all from the *Veillonellaceae* family which is likely the explanation for their correlation. Members of the *Veillonellaceae* family have been linked with inflammation [64].

By increasing statistical power and providing context for the relationships between significant taxa, SCNIC modules open new opportunities for analysis. When a module is associated with a variable of interest, the correlations within the module may imply functional relationships. These can be further investigated with *in vitro* and *in vivo* experiments. Studies which aim to test hypotheses generated from correlative analysis will commonly use a single significantly associated microbes. This often does not adequately represent *in vivo* systems because microbes in isolation often do not affect a disease state or their environment. SCNIC can enhance these confirmatory studies by identifying groups of microbes that may grow better than individual microbes and may better elicit relevant phenotypes than when grown separately.

## ACKNOWLEDGEMENTS

We would like to thank Elmar Pruesse for input on the design of SCNIC. We thank Jennifer Fouquier, Abigail Armstrong and Casey Martin for beta testing SCNIC. We thank Jennifer Fouquier for reading and commenting on the manuscript. Funding for KT came from the University of Colorado School of Medicine Research Track. Funding for MS came from NIH NLM 4 T15 LM009451-10.

## DATA ACCESSIBILITY

The Noguera-Julian et al. data set is available from NCBI SRA accession number SRP068240. The lakes dataset is available from QIITA accession number 1041.

## AUTHOR CONTRIBUTIONS

M.S. coded the initial implementation of SCNIC and made major contributions to its conceptualization and design. K.T. improved the SCNIC implementation and performed the HIV case study. J.S. performed the comparisons of SMD to LMM with real and simulated data. C.L. conceptualized SCNIC and guided its implementation and design. K.T., M.S., J.S. and C.L. all wrote the manuscript together.

## Lakes Datasets

ANCOM analysis of the HIV and Great Lakes datasets after using SCNIC at R-Value thresholds at 0.2, 0.4, and 0.65. MSM was used as the categorical variable for differential abundance in the HIV analysis and the Great Lakes analysis tested for taxa that differed between Lakes.

**Supplemental Table 1:** Runtime and memory usage by SCNIC module generation step. Modules were calculated based on correlation matrices for the Great Lakes dataset and an integrated metabolomics-microbiome (GT) dataset. Runtime was profiled using GNU Time across 3 runs per dataset and method, and maximum memory usage in mebibytes (MiB) was profiled using memory-profiler. A count table to sum module relative abundances was passed to SCNIC for the GT dataset when denoted in the leftmost column.

| Dataset | Features | Time 1 | Time 2 | Time 3 | Max Memory |
| --- | --- | --- | --- | --- | --- |
| Lakes (SMD) | 764 | 0 m 31 s | 0 m 30 s | 0 m 32 s | 190 MiB |
| Lakes (Louvain) | 764 | 0 m 7 s | 0 m 7 s | 0 m 7 s | 190 MiB |
| GT (SMD) | 1,301 | 1 m 28 s | 1 m 27 s | 1 m 27 s | 302 MiB |
| GT (SMD + table) | 1,301 | 1 m 45 s | 1 m 32 s | 1 m 52 s | 301 MiB |
| GT (Louvain) | 1,301 | 0 m 20 s | 0 m 18 s | 0 m 16 s | 299 MiB |
| GT (Louvain + table) | 1,301 | 0 m 15 s | 0 m 15 s | 0 m 16 s | 301 MiB |

